# The Role of Resident Custodian Communities in Devolved Forest Conservation: Lessons from a retrospective, mixed-method observational study of the Ifakara-Lupiro-Mang’ula Wildlife Management Area in Southern Tanzania

**DOI:** 10.1101/2025.10.20.683348

**Authors:** Lucia J. Tarimo, Deogratius R. Kavishe, Mohamedi Kamuna, Jackline Nyaruzi, Clarence Msafiri, Fidelma Butler, Felister Mombo, Gerry F. Killeen

## Abstract

The establishment of Wildlife Management Areas (WMAs) in Tanzania aims to involve local communities in conserving natural resources on their land, while also yielding sustainable socio-economic benefits. WMAs are established on community lands to ensure the protection of wildlife and their habitats along the fringes of centrally managed protected areas like National Parks and Game Reserves. A recent cross-sectional study of the Ifakara-Lupiro-Mang’ula (ILUMA) WMA in the south of the country suggests that small, authorized fishing camps established legally inside the designated conservation area act as resident custodian communities who actively collaborate with the WMA to enforce conservation of the surrounding forest ecosystem. Here a retrospective cartographic analysis of a longitudinal series of surveys of human, livestock and wildlife activities across ILUMA between 2022 and 2025 reveal a clear increase of illegal timber harvesting in forested areas around custodian communities immediately after they were displaced by extreme floods in 2024. Furthermore, recorded narratives of resident members of these fishing camps indicate that charcoal burners, timber harvesters and even game meat poachers have taken advantage of such absences to exploit the forest undetected. These observations further support the view that resident custodian communities living within WMAs may play vital roles in conservation efforts, particularly in settings where financial resources for enforcement of regulations are limited.

## Introduction

Community based natural resources management (CBNRM) models have been widely implemented across Africa for over four decades, prompted by the well-documented failure of “fortress conservation” in many contexts (Fortmann et al., 2001; Gibson & Marks, 1995; Nelson & Agrawal, 2008; Songorwa, 1999). These models aim at ensuring sustainable utilization and conservation of natural resources through active participation and empowerment of local communities. The overall goal of CBNRM is to achieve balance between conservation with socio-economic development priorities, thereby ensuring that communities benefit from the sustainable use of resources like wildlife and forests, while also preserving them for the future generations (Gibson & Marks, 1995; Goldman, 2003). The underlying assumption of CBNRM is that communities are more likely to successfully engage in conservation efforts when they are given ownership, authority and responsibility for managing their own local natural resources (Agrawal & Gibson, 1999; Kull, 2002; Murphree, 2004). Furthermore, it also assumes that the benefits local communities can derive from sustainable resources use, such as consumptive and non-consumptive tourism activities, will motivate them to protect and manage their environment effectively (Murphree, 2004). However, these models have achieved mixed outcomes in practice, with many documented success and failures (Balint & Mashinya, 2006; Salerno et al., 2016; Shackleton & Campbell, 2000; USAID, 2013).

In Tanzania, an important mechanism through which the wildlife sector practices decentralized CBNRM is a legally formalized institutional model known as Wildlife Management Areas (WMA) (Government of Tanzania, 1998). These community-owned local organizations collectively manage consolidated areas of village land that contain ecologically significant habitats that are rich in wildlife. An important aim of establishing such devolved WMAs is to ensure the protection of these natural resources along the fringes of centrally managed protected areas like National Parks and Game Reserves, while also providing sustainable socio-economic benefits to the stakeholder communities who set aside land for their establishment (Baldus & Cauldwell, 2004; Government of Tanzania, 1998). The formal implementation of the WMA governance model started in 2003 and there were 24 WMAs established across the country that have been gazetted and granted user rights at the time of writing (Government of Tanzania, 2023). Each of these areas operates under its own set of locally-tailored bylaws that complement the national WMA regulations and the Wildlife Policy (Government of Tanzania, 1998, 2012). The stakeholder villages actively establish an Authorized Association (AA) that contributes to the formulation of these local bylaws, to optimize effective and sustainable utilization of relevant natural resources. However, the effectiveness of these WMAs varies, as some are struggling to meet their conservation and community development objectives, because of challenges arising from human encroachment, inadequate income and limited management capacity (Bluwstein et al., 2016, 2018; Keane et al., 2020; Kimario et al., 2020; Mwakaje, 2010; USAID, 2013, 2016).

One recently studied example in the south of the country is the Ifakara-Lupiro-Mang’ula (ILUMA) WMA (Figure 1). This WMA is challenged by increasing human encroachment, making it difficult to achieve both its conservation and socio-economic development objectives (Duggan et al., 2025; Tarimo et al., 2025). However, recent studies conducted across the area provide evidence of high ecosystem integrity in parts of the WMA where small resident communities are conditionally allowed to live within the conservation area and engage in regulated fishing activities for livelihood purposes (Duggan et al., in press). The presence of these resident fishing camps appears to contribute to conservation of their surroundings through collaboration with the management of the WMA, effectively providing year-round community-based surveillance for unauthorized activities like charcoal burning, timber harvesting, meat poaching, cattle herding and illegal fishing practices. As such, they may be described as *custodian communities*, whose continuous presence as active participants in WMA conservation activities represents a notable exception to the general rule that human settlement is closely associated with disturbance of natural ecosystem (Duggan et al., 2025, in press).

**Figure 1.**
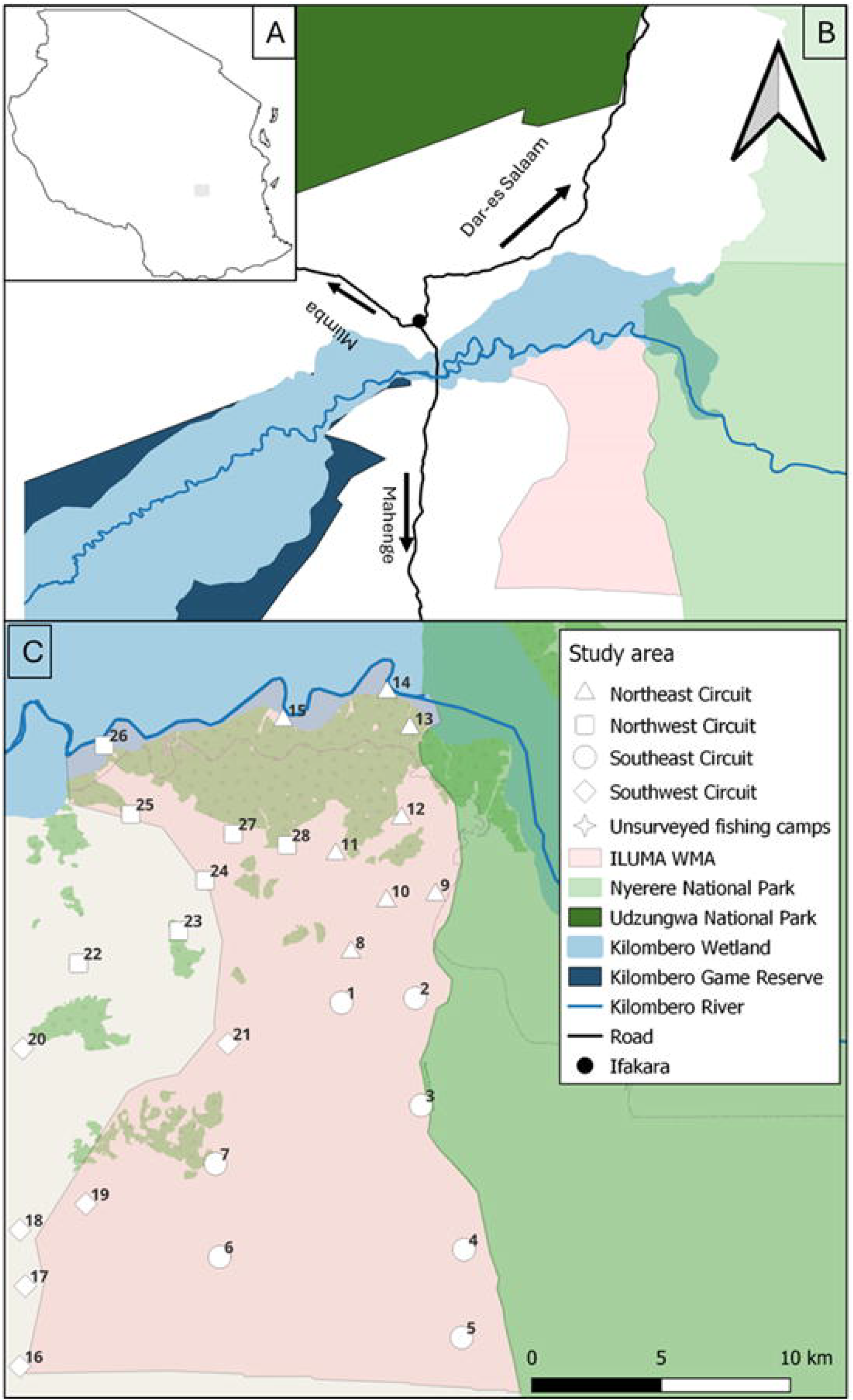
Map displaying the surveyed area in and around the Ifakara-Lupiro-Mang’ule Wildlife Management Area (ILUMA WMA), framed in a local and national context in southern Tanzania. The insert at the top left is the map of Tanzania (**Panel A**), with the pinkish-coloured area representing the designated conserved lands of the Ifakara-Lupiro-Mang’ula ILUMA to the south of Kilombero River **(Panel B**). To the East of ILUMA is Nyerere National Park, while to the west lies the extensive wetlands of the Kilombero Valley inland delta and Ifakara town to the north-west. All the visited locations where the survey team usually stayed overnight (Duggan et al., 2024, 2025, in press) are referred to as camps and illustrated in **Panel C**. Twenty-two of these camps are within the protected conservation area of the ILUMA WMA, while six are in or near the adjacent villages just outside it immediately to the west. As described in section 2.3, surveys of human, livestock and wildlife activities were conducted within a 2km radius around these camp locations (Radial surveys) and while hiking from one camp to the next (Transect surveys) (Duggan et al., 2024, 2025, in press).

While the cross-sectional evaluation of Duggan et al. (2025, in press) revealed robust natural ecosystem integrity in the forested areas immediately around these authorized fishing camps, here we report a mixed method study providing further retrospective longitudinal evidence for their invaluable roles as custodian communities. Specifically, a retrospective analysis of a longitudinal series of human, livestock and wildlife activities surveys is combined with shared recollections of community stakeholders, to illustrate how illegal harvesting of timber, charcoal and game meat became more frequent around these authorized settlements during periods when they were abandoned because of extreme floods or an enforced ban on fishing in the area.

## 2. Methods

### 2.1 Study area

This study was conducted in the ILUMA WMA, in the Kilombero Valley of southern Tanzania, the largest lowland freshwater wetland in east Africa and a recognized RAMSAR site (Dinesen, 2016). The conserved area of the WMA encompasses 509km^2^ of village lands, spanning both the Kilombero and Ulanga Districts in Morogoro region of Southern Tanzania. To the east, it borders Nyerere National Park, which is the largest national park in Africa. The ILUMA WMA features a variety of vegetation cover, including *miombo* woodland, dense groundwater forest, corridors of riparian forest along water courses, open grasslands and wetlands. It is also home to diverse range of wild animals including elephant, buffalo, leopard and lion (Duggan et al., 2025, 2024).

The area experiences a short rainy season, characterized by sporadic but sometimes intense rain occurring between October and December, while the long rainy season is more predictable and consistent, occurring between March and May. Between June and October, the area experiences little to no rain. The centre of the valley lies on the attitudes of approximately 300m with annual mean daily average temperature of 24°C (Sigalla et al., 2023).

The WMA consists of 15 stakeholder villages (Tarimo et al., 2025), who have a mandate from central and local government to decide how they can use their resources sustainably, so long as they do so within the constraints defined by the Tanzania Wildlife Management Area Regulations and the Wildlife Policy (Government of Tanzania, 1998, 2012). The Kilombero River flows through the north of the ILUMA, where the surrounding communities are legally allowed to engage in regulated fishing activities and reside at well-defined, pre-agreed locations within the conservation area, under the condition that they also cooperate with the WMA authorities, thereby acting as resident custodian community (Duggan et al., in press).

### 2.2 Study design

This study is a retrospective, mixed-method observational design, which used data from a longitudinal series of surveys of human, livestock and wildlife activities across the ILUMA WMA and adjacent settled areas along its western boundary. The sampling framework used for surveying the area consists of 28 different locations referred as camps (Figure 1), the details of which have been described previously (Duggan et al., 2024, 2025, in press). Twenty-two of these camps are within the protected conservation area of the ILUMA WMA, while six are outside in or near the adjacent villages immediately to the west. It should be noted that this study was carried out as part of a larger project related to malaria vector mosquito ecology (Kavishe et al., 2024; Walsh et al., in press), which determined the selection of these locations for radial surveys around surface water bodies within a 2km of each camp and, consequently, the routes taken between them for transect surveys (Duggan et al., 2024, 2025, in press).

The 22 camps within ILUMA were distributed across the entire area to ensure coverage of a variety of different environments. These included areas with heavy human activity, open and dense miombo woodland, dense groundwater forest and three of the five authorized fishing camps at the edge of the large dense groundwater forest along the southern bank of Kilombero River. The 6 camps just outside the western boundary were all either close to established substantial villages or among the scattered farms of more dispersed rural communities. Data were collected in seven distinct rounds of surveys between January 2022 and July 2025, initially spanning different seasons of the year in 2022 and 2023 but then being repeated annually in the early to mid-dry season in 2024 and 2025. Not every camp was visited in each round due to practical circumstances in the field e.g. flooding in the rains or lack of surface water in the dry season (Duggan, 2023) but the majority of them were visited in all rounds.

### 2.3 Field surveys of human, livestock and wildlife activities

Data were collected using two methods: radial surveys around surface water bodies within a 2km radius of each camp and line transect surveys undertaken while hiking from each camp to the next (Duggan et al., 2025, 2024, 2025 in press). The data collected included both direct and indirect observations of the activities of humans, livestock and wildlife. While a wide range of indicators of different human activities, as well as various signs of activity by livestock and wildlife were recorded, the only indicators assessed and presented herein relate to unauthorized utilization of forestry resources, specifically timber harvesting and charcoal burning. The field-estimated age of each sign of these activities was also recorded, with each observation classified as either active, recent, or old. In this age classification system *active* referred to activities that appeared to have occurred within the same season as it was observed, *recent* to those from the previous season, and *old* to those from any season before the preceding one.

In each camp, GPS coordinates were recorded, and then radial surveys were conducted following along the fringes of any surface water body like puddles, streams, waterholes, rice fields or rivers within the area immediately surrounding each camp. This procedure was designed to align with the protocol followed by a team surveying mosquito larvae (Walsh et al., 2025 in press). The area covered by these radial surveys was 2km radius in any direction from the camp and sampling was limited to a maximum four-hour time frame, to define practical limits to the survey duration (Walsh et al., 2025 in press).

The line transect surveys were conducted on foot while hiking from one camp to the next with groups of 6 to 8 camps that were arranged along circuits that could be visited in sequence, each corresponding approximately to a single northeast, northwest, southeast or southwest quadrant of the study area (Duggan et al., 2024, 2025, in press). The starting point for each transect was the camp being departed from, and each transect was divided into a series of up to 10 transect segments 500m long, using a Garmin® eTrex GPS to define where each segment started and ended. Throughout each 500 m segment, all relevant observations of human, livestock or wildlife activities were recorded using a standardized set of data interpretation and recording tools (Duggan et al., 2024, 2025, in press). The number of segments surveyed along each route between camps depended on the distance between the camps. If the distance was less than 5 km, transect segments were recorded until the new camp was reached. If exceeded 5 km, a maximum of 10 segments were completed, after which the survey was stopped, and the remaining distance was completed without collecting data. This limit was set to ensure that data collection could be carried out with consistent quality and full concentration without being compromised by investigator fatigue.

### 2.4 Descriptive stakeholder recollections

Additional qualitative sociological data, in the form of publicly expressed retrospective perspectives of relevance to this study, were collated by the lead investigator during informal conversations with stakeholder communities over the course of the carrying out the activity surveys described above and also at a stakeholder workshop she attended. The perspectives considered of relevance to this study were extracted from the field notes of the lead investigator, who wrote down what was said verbatim. The workshop was organized by the African Wildlife Foundation (AWF) with the purpose of electing *Conservation Champions* for the Kilombero Valley. Stakeholders from the ILUMA WMA and representatives from the fishing camps in the WMA attended the workshop to take part in the process. Some of the perspectives shared openly by stakeholders at this meeting were relevant to and consistent with the observations of the investigators, so they were noted and written down at the time by the lead author. All quotes presented herein were originally recorded as hand-written notes in *kiSwahili* and then translated into English to be reporting herein.

### 2.5 Data Presentation

The entire dataset of human, livestock, and wildlife activities for seven rounds was subset in *RStudio*® to focus specifically on active charcoal burning and timber harvesting activities from both radial and line transect surveys. To show the number of observations of these activities around each camp in radial surveys and between camps along transect survey segments, maps were created in *QGIS*® to cartographically represent their distribution across the ILUMA WMA. This was accomplished by adjusting the size of the symbols used to represent each surveyed camp or transect segment on the maps, with the magnitude of each one representing the number of observations for each particular activity category.

## 3. Results

Despite the ready availability of high-quality timber within the groundwater forest along the south bank of the Kilombero river, illegal harvesting was rarely encountered in this northernmost part of the WMA over the first five rounds of activity surveys that spanned the period from January 2022 to October 2023 (Figure 2). On the sixth round of surveys in mid-2024, however, it became widespread across that groundwater forest, before again becoming conspicuous by its absence in the seventh round in mid-2025 (Figure 2). Timber harvesting was never encountered within that essentially intact groundwater forest during rounds 1, 3 or 7, only once during rounds 2 and 4 and four times on round 5. In stark contrast, active timber harvesting was observed 13 times in round six, which immediately followed exceptional floods that displaced two of the surveyed fishing camps and two more unsurveyed ones. Remarkably, four occurrences of active timber harvesting were observed in the displaced fishing camp of Funga (Camp 26) and 6 times close to the almost completely displaced fishing camp of Mdalangwila (Camp 15), with two more at the edge of the forest *en route* to the latter. By comparison, only one observation of active timber harvesting was made in the sixth survey round near the fishing camp of Mikeregembe (camp 14), which is built on a high bank of the river and was largely unaffected by floods. Although active timber harvesting was frequently observed elsewhere across the WMA throughout the study period, its distribution was scattered and sporadic, varying unpredictably from one survey to the next. Outside the large groundwater forest along the south bank of the Kilombero River, no active timber harvesting was observed inside the protected area of the WMA during rounds 1,3, 4 or 7, while four occurrences were observed in round 2, six in round 5 and nine in round 6.

**Figure 2:**
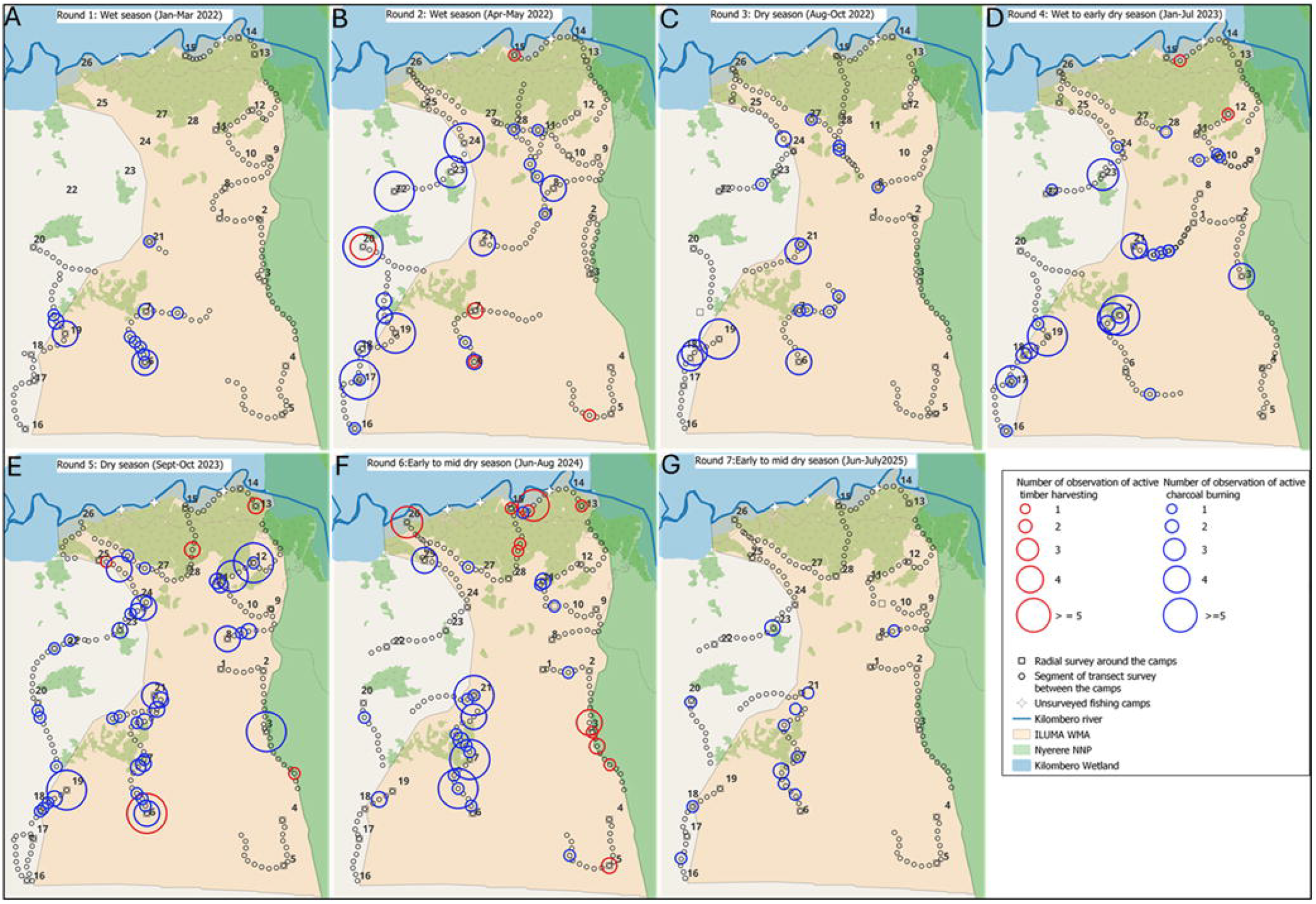
Maps showing active timber harvesting and charcoal burning detections across the Ifakara-Lupiro-Mang’ula Wildlife Management Area (ILUMA WMA) over activity survey rounds 1 to 7. The authorised fishing camps in the north of the WMA are camp numbers 14, 15 and 26 along the banks of the Kilombero River. The size of the red and blue circle symbols respectively indicate the number of recorded observations of active timber harvesting and charcoal burning around each camp or along each transect segment.

By comparison, charcoal burning was much more common across the WMA, except for the groundwater forest around the fishing camps (Figure 2). Out of a total of 263 observations of active charcoal burning over the first five rounds of activity surveys, it is remarkable that none occurred in this heavily forested northernmost area near the Kilombero River. Notably, the only observation of active charcoal burning in the northern groundwater forest area was made in the sixth round of activity surveys, immediately after two of the surveyed fishing camps had been displaced by floods. Indeed, it was particularly remarkable that this only instance of active charcoal burning occurred right beside the normally busy pathway that encroachers usually avoid because it is frequently used by residents of these custodian communities and the traders who regularly visit them. In comparison, no active charcoal burning was recorded around this northern groundwater forest area during the seventh round, which was conducted after the custodian communities returned.

We initially learned about the displacement of these custodian communities by severe flooding between March and May 2024 while visiting and staying overnight at these fishing camps during the sixth round of activity surveys. The informal narratives shared by community members then, and during similar visits during the seventh round of activity surveys, indicated charcoal burners and timber harvesters took full advantage of that opportunity to enter the forest unobserved. This was further noted during the Kilombero Valley Conservation Champions workshop organized by AWF, at which one participant said the following:

Quote 1: *“Even in this year when there were heavy floods, we moved from our camps because water spread along. … At this time, charcoal burner and timber harvesters have badly destroyed our forest*.” Fishing Camps Representative, Kilombero Valley Conservation Champion’s Workshop, August 2024.

Members of these custodian communities further explained that timber harvesters often enter in the forest when they are absent, especially during the night. It was also obvious at the workshop that these custodian community representatives understood their active roles in the conservation of the area and the impact that their participation has. Specifically, they explained that they also noted decreases in wild animal activity associated with unauthorized deforestation activities that increased in their absence, notably during a historical period of absence much longer ago due to a temporary ban on fishing:

Quote 2: *“We contribute a lot to conserving our forest. We report any illegal activity or anyone entering the forest without a permit. We also ensure that unauthorized people do not enter… I remember when fishing activities were closed from November 2018 to June 2019. When we returned, we did not see wildlife as frequently as before. In our absence, poachers took advantage and entered the area because there was no one to report them*.*”* Fishing camp Representative, Kilombero Valley Conservation Champion’s Workshop, August 2024.

## 4. Discussion

The findings of this study are consistent with evidence that community participation can enhance conservation efforts (Anagnostou et al., 2020; Bhatta et al., 2018; Gaodirelwe et al., 2020; Lee, 2018; Lee & Bond, 2018; Pandey et al., 2014; Steinmetz et al., 2014). The various positive outcomes attributed to community engagement in conservation have consistently been associated with increased sense of ownership over their natural resources and tangible benefits accrued from conservation. Specifically, this study adds to that evidence base by illustrating how legally permitted, carefully managed communities of fishers residing within the ILUMA WMA, with authorized and exclusive access to fish resources can serve as custodians of the surrounding land. In this specific case, the residents of these fishing camps appear to play a crucial role in protecting the most densely forested part of ILUMA, through the collaboration with the WMA management committee and Village Game Scouts. This has also been noted in MBOMIPA WMA, where community members actively participated in reporting of illegal activities to the WMA management, thereby contributing positively to conservation of the area (Mdete, 2016).

Previous cross-sectional surveys of the ILUMA WMA indicated that these forested parts near these authorized fishing camps had higher ecosystem integrity than other inhabited parts of the WMA (Duggan et al., 2025, 2025 in press). These findings by Duggan et al. (2025, in press) are further reinforced by the retrospective observations reported herein, illustrating how the displacement of these custodian communities by flooding enabled unmonitored deforestation through illegal harvesting of timber and charcoal. Therefore, we recommend that future research should seek more in-depth insights into how these custodian communities function and how they are motivated, to better understand the principles underpinning this success story.

This study also sheds light on the relative value of different forestry resources and informs best practice for conservation of such areas looking ahead. Although a few instances of timber harvesting were detected in the ground water forest in some of survey rounds, even when the fishing camp custodian communities were present, charcoal burning was never observed even once under such circumstances. The most obvious interpretation of this observation is that it is common for timber harvesters to take greater risks because the higher economic value of hardwood timber than charcoal. Additionally, the single detection of charcoal burning observed in the survey conducted immediately after the displacement of the custodian communities, which was never observed before or afterwards, even that single observation may be of significance. The fact that this singular instance of charcoal burning was so brazenly carried out along a main access route that is used so frequently throughout most years raises the concern that charcoal burning could rapidly increase within this forest if these custodian communities were closed without the WMA implementing more frequent patrols to compensate for the loss of their roles in conservation monitoring.

Like all studies, this one has considerable limitations, the most important of which is that simply it is purely observational in nature. This opportunistic retrospective study was prompted by fortuitous observations made after the unforeseen displacement of custodian communities from authorized fishing camps by extreme floods. It also relied on the existing data from a longitudinal study that used radial and line transect surveys, respectively conducted around convenient locations for overnight camping and trails for hiking on foot between them. However, this survey framework may not have adequately covered large areas that are densely vegetated and less accessible, particularly those deep within the forest surrounding these communities. Future research should further explore the long-term effectiveness of custodian communities in conservation activities and consider similar approaches to alternative livelihoods for resident custodian communities that are based on the sustainable utilization of natural resources other than fish, to better cover larger areas away from major river.

With livestock keeping also identified as a major challenge within the ILUMA WMA (Tarimo et al., 2025), the ILUMA community could develop solutions to ensure the availability of livestock fodder through silage production (Muck et al., 2018; Tisocco et al., 2024; Wang, 2025) from the excess of tall grasses grow in the numerous glades distributed across that southern miombo woodland area of the WMA. Such selective seasonal harvesting of excess grass to provide affordable fodder to neighbouring pastoralist communities all year round could help to reduce the pressure of livestock grazing within the conservation area, especially during the rainy season when fertile land in this area is typically dedicated to crop cultivation (Duggan et al., 2025, in press). Moreover, having resident custodian communities producing silage within these areas could also enable passive monitoring of encroachment at negligible cost, rather than exclusively relying on active patrol by VGS who need to be paid for such visits.

## Supporting information

https://doi.org/10.5281/zenodo.15161027

## Acknowledgement

The authors wish to thank the Village Game Scouts of ILUMA WMA for their hard work and enthusiastic participation in field activities. We also thank all the governance, management and community stakeholders in the ILUMA WMA for all their kind collaboration and assistance. We express our appreciation to Mr Rogath Msoffe at the Ifakara Health Institute, Mr Arstid Litaka, and Mr Patric Cyprian from Sokoine University of Agriculture, and Ms Lily Duggan, Ms Katrina Walsh, Ms Sarah Cheallaigh, Ms Yasmeen Hadjieva and Mr Jude O Regan from University College Cork for their participation in the activity surveys. We also thank Dr. Emmanuel Kaindoa, Mr. Frederic Masanja, and Mr. Fadhili Songo for the essential institutional support provided by the Ifakara Health Institute throughout the course of the study. We would also like to thank Dr. Ronan Hennessy, Dr. Ramiro Crego and Dr. Tom Reed at the University College Cork for their kind help with analytical skills training. A very special word of thanks is due to our recently deceased friend and colleague, Mr Octavian Malopola, without whom this work would never have even begun.

## Authors contribution

Lucia J. Tarimo: Conceptualization, investigation, methodology, implementation, formal analysis, writing original draft of the manuscript. Deogratius R. Kavishe: Methodology, review, editing, validation. Mohamedi Kamuna Review, editing and validation. Clarence Msafiri: Methodology, review, editing and validation. Jackline Nyaruzi: Methodology, review, editing and validation. Fidelma Butler: Conceptualization, review, editing, validation. Felister Mombo: Conceptualization, review, editing and validation. Gerry F. Killeen: Acquisition of fund, conceptualization, methodology, implementation, review, editing and validation.

## Data availability statement

All data are fully available without restriction

## Supporting information

**Supplementary file S1 Data:** Seven round survey data on the observations of active charcoal burning and timber harvesting across ILUMA WMA.

